# Intestinal helminth infection transforms the CD4+ T cell composition of the skin

**DOI:** 10.1101/2021.04.18.440186

**Authors:** C H Classon, M Li, J Ma, A Lerma Clavero, X Feng, C A Tibbitt, J M Stark, R Cardoso, E Ringqvist, L Boon, E J Villablanca, A Gigliotti Rothfuchs, L Eidsmo, J M Coquet, S Nylén

## Abstract

Intestinal helminth parasites can alter immune responses to vaccines, other infections, allergens and autoantigens, indicating effects on host immune responses in distal barrier tissues. We herein show that C57BL/6 mice infected with the strictly intestinal nematode *Heligmosomoides polygyrus* have impaired capacity to initiate skin immune responses and develop skin-resident memory cells to mycobacterial antigens, both during infection and months after deworming therapy. Surprisingly, and in contrast to a previously noted loss of T cells in peripheral lymph nodes, the skin of worm-infected mice harboured higher numbers of CD4+ T cells compared to skin of uninfected controls. *H. polygyrus-*specific T_H_2 cells accumulated during infection and remained after worm expulsion. Accumulation of T_H_2 cells in the skin was associated with increased expression of the skin-homing chemokine receptors CCR4 and CCR10 on CD4+ T cells in blood and mesenteric lymph nodes draining intestinal tissues, indicating gut-to-skin trafficking of cells. In conclusion, we show that infection by a strictly intestinal helminth has long-term effects on immune cell composition and local immune responses to unrelated antigens in the skin, revealing a novel mechanism for T cell colonization and worm-mediated immunosuppression in this organ.

## Introduction

Immune responses in the gut and skin are deeply intertwined, as indicated by skin manifestations of intestinal disorders such as inflammatory bowel disease (IBD), Coeliac disease, small intestinal bacterial overgrowth and food allergies. ^1^ The reduced prevalence of helminth infections in modern times has been linked to an increase in autoimmune and inflammatory disorders in Western societies. 2, 3 Intestinal worm infections have been suggested to have beneficial effects on inflammatory skin disorders with reduced atopy and allergic reactions in the skin. ^2, 3^ We and others have shown that worm infection can dampen immune responses to infection and vaccination in the skin. ^4-6^ Enhanced T helper-cell type 2 (T_H_2) and regulatory (T_REG_) responses are commonly suggested to drive the down-modulating effects of worms on subsequent immune responses. ^2^ Furthermore, we recently described how a chronic intestinal infection with the model worm pathogen *Heligmosomoides polygyrus bakerii* causes redistribution of circulating lymphocytes with accumulation of T cells in the mesenteric lymph nodes (mesLNs). ^6^ Worm-induced expansion of mesLNs resulted in atrophy and loss of T cells in skin-draining lymph nodes (LNs), leading to dampened responses in the skin to the tuberculosis vaccine *Mycobacterium bovis* Bacillus Calmette Guérin (BCG). ^6^

Tissue-resident T cells (T_RMs_) in barrier tissues contribute to infection- or vaccine-driven immunity against pathogens and also to inflammatory disorders. ^7-11^ Given the dampening effect of worms on skin inflammation and the importance of skin T_RMs_ in local immune responses, we here assessed the impact of intestinal infection with *H. polygyrus* on skin T-cell composition and function. *H. polygyrus* infects orally through intake of L3 larvae and distributes strictly to the gut where it establishes a persistent, chronic infection. ^2^ The responses to *H. polygyrus* in the local intestinal mucosa are well described. ^2, 6, 12^ Alike other intestinal worm infections, *H. polygyrus* induces a T_H_2 response causing goblet cell hyperplasia and increased peristaltic movements that provoke worm expulsion. In the chronic phase of *H. polygyrus* infection, T_REG_ responses dominate and ameliorate the pathological immune responses. ^2, 6, 12^ As mice infected with *H. polygyrus* lose lymphocytes from skin-draining LNs ^6^ we hypothesized that intestinal worms would also affect the lymphocyte composition in the skin. Accordingly, we found that *in situ* recall responses to mycobacterial lysate were reduced in animals with chronic *H. polygyrus* infection. In contrast to T cell loss in skin-draining LNs, *H. polygyrus* promoted deposition of worm-specific T_H_2 cells in the skin. Interestingly, these T cells remained in the skin for months after worm expulsion. Our findings show that persistent parasitic infections imprint distal host tissues with T_H_2 cells. Importantly, we propose that skin-deposited T_H_2 cells contribute to the altered skin immune response upon subsequent pathogen challenge or intradermal delivery of vaccines, providing a novel mechanism to previously observed immune-regulations in skin of worm-infected mice.

## Results

### Intestinal H. polygyrus infection dampens skin recall responses to mycobacterial antigens

We have previously shown that mice infected with *H. polygyrus* mount weaker delayed type hypersensitivity (DTH) responses to BCG. ^6, 13^ To investigate effects of intestinal worms on development of mycobacteria-specific memory T cells in the skin, local cytokine recall responses were assessed 10 weeks after injection of whole cells lysate (WCL) from *M. tuberculosis* in ear skin of worm-infected mice (Fig. 1a). WCL was used to avoid confounding factors associated with live mycobacterial challenges, such as microbial spread and replication. The long interval between WCL injection and recall assessment was used so that WCL-induced inflammation and effector responses would have waned. Ear thickness was measured at various time points after WCL administration, but no significant difference was observed between worm-infected and worm-free mice (Supplementary Fig. 1a). As ear thickness was not fully restored but rather stabilized after 8-10 weeks, we also analysed the contralateral, untouched ear. Interestingly, recall IFN-γ production in WCL-injected and untreated ears were lower in *H. polygyrus-*infected mice compared to worm-free controls (Fig. 1b, 1c, 1d, 1f, g, Supplementary Fig. 1b, 1c). Surprisingly, CD4+ T cell numbers were higher in ears of *H. polygyrus-*infected mice in both WCL-treated and untreated ears (Fig. 1e, h). Hence, mice infected with *H. polygyrus* mount weaker skin recall responses to mycobacterial WCL both at the site of injection and as well as distal skin sites, despite an accumulation of CD4+ cells in the skin.

**Figure 1:**
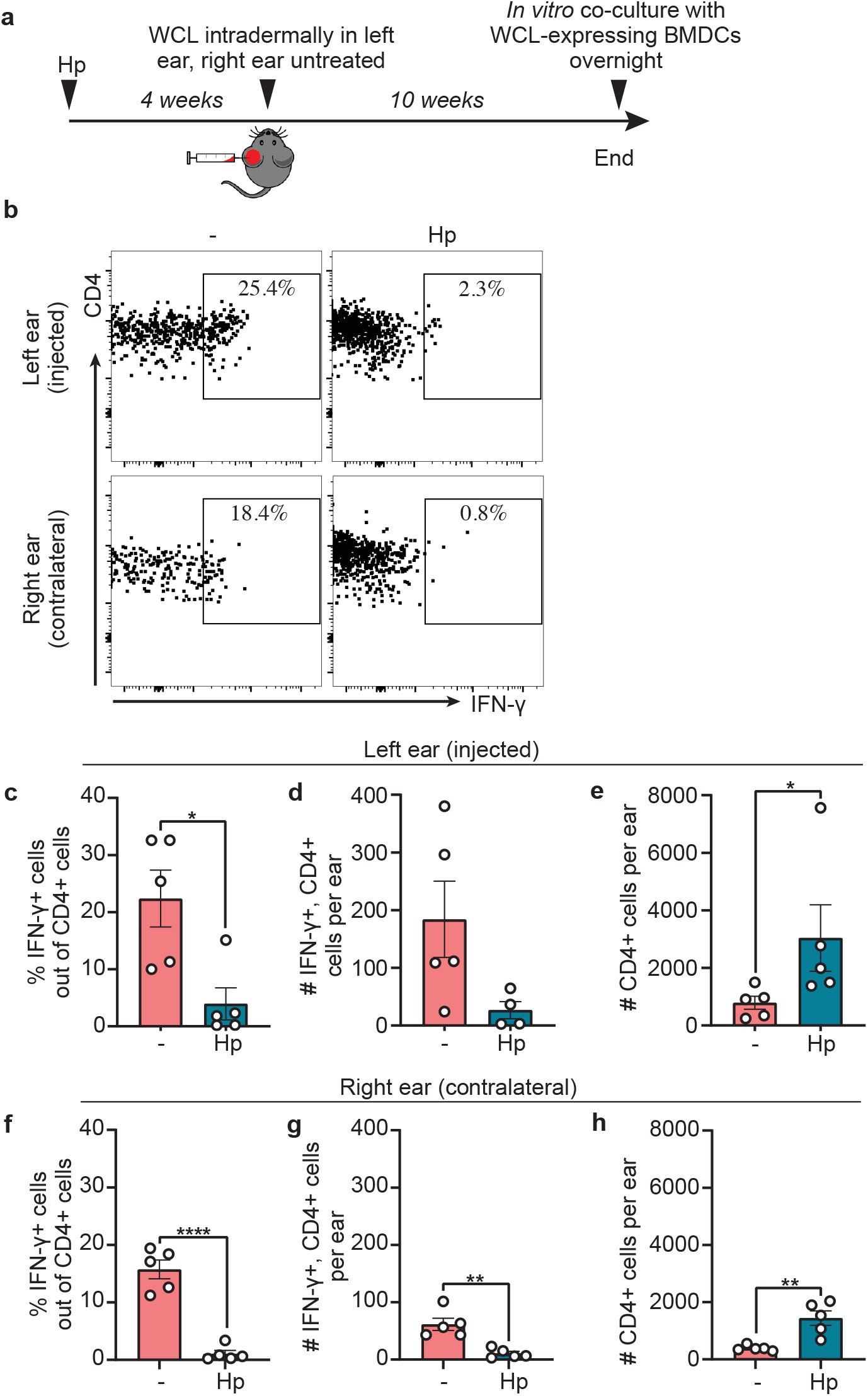
Intestinal *H. polygyru*s infection dampens recall responses to mycobacterial antigens in the skin. Mice were infected with *H. polygyrus* (Hp) or not (-) and treated according to experimental outline **a**: Four weeks after Hp infection, one ear was injected intradermally with whole cell lysate (WCL) from *Mycobacterium tuberculosis*, and the contralateral ear left untouched. Ten weeks later, ear cells were cultured with BMDCs expressing WCL overnight and analysed by flow cytometry. **b** Representative FACS plots illustrating the gating strategy used to identify IFN-γ+ CD4+ cells in injected (**c-e**) and contralateral, untouched (**f-h**) ear. One out of two independent experiments with similar results are shown. Each dot represents one mouse (n=5) and bars indicate mean ± SEM. Statistical differences are depicted as *p < 0.05, **p < 0.01, and ****p < 0.0001.

### Intestinal H. polygyrus infection, but not DSS-induced colitis, causes T cells to accumulate in skin

To further explore how *H. polygyrus* infection affects skin T cell responses, we analysed cells in the skin of mice after worm infection without further manipulation. Succeeding experiments were performed on back skin, providing a larger surface area. Two weeks after *H. polygyrus* infection, skin CD4+ T cell numbers began to increase (Fig. 2a, 2b, Supplementary Fig. 2a) and by 4 weeks after infection T cell numbers were 2-3 times higher than in the skin of uninfected mice (Fig. 2c-2d). No differences were noted in CD8+ cell numbers at either time point (Fig. 2a-2d). Most CD4+ T cells in skin of both worm-infected and worm-free mice co-expressed CD44 and CD69, indicating an effector or memory phenotype (Fig. 2e-2f).

**Figure 2:**
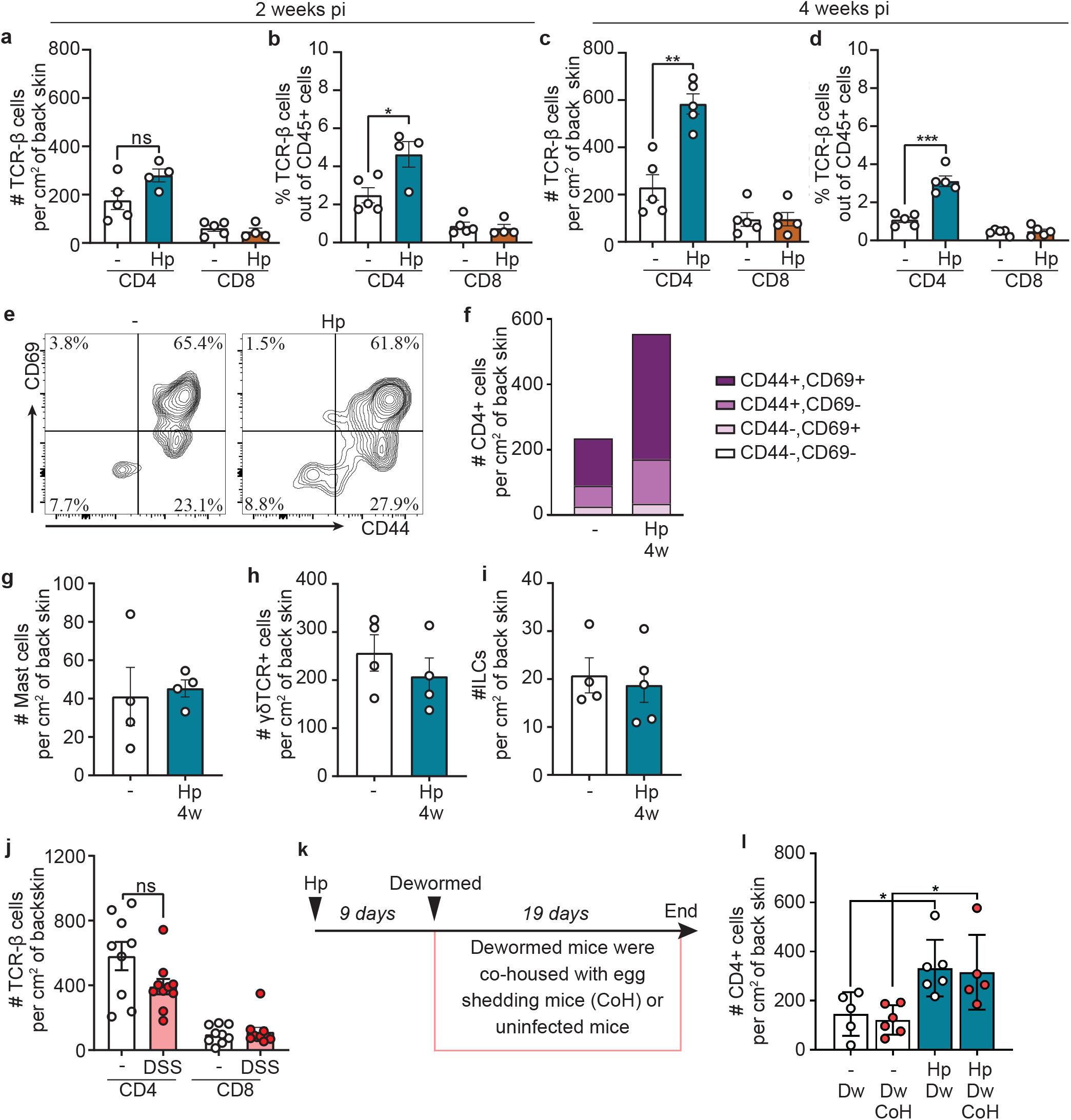
Intestinal *H. polygyrus* infection, but not DSS colitis, causes T cells to accumulate in skin. **a-i, l** Mice were infected with *H. polygyrus* (Hp) or not (-). **a-d** Absolute numbers and frequencies of CD4+ and CD8+ T cells found in back skin 2 and 4 weeks post infection (pi). **e** Representative FACS plots of CD44 and CD69 gating. **f** Absolute numbers and percentages of CD44 and CD69 out of CD4+ T cells. Data is calculated from an average of three independent experiments with a total of ≥ 12 mice per group. **g** Absolute numbers of mast cell (c-Kit+, FcεRI+, linage-(lin-) cells), **h** γδ TCR+ cells and **i** ILCS (CD45+, lin-, CD90.2+). **j** Mice were treated for 2 x 7 days with 2% dextran sodium sulfate (DSS) in the drinking water with two weeks in between the treatments. **k** Experimental outline for **l**: De-wormed mice were co-housed with either uninfected or infected (CoH), egg-shedding mice. Egg-shedding mice were infected with 500 L3 Hp to ensure high egg production. **a-e, g-i, l** One out of at least two independent experiments with similar results are shown. In **j**, the experiment has been performed once but with 10 mice per group. Each dot represents one mouse (n≥4, **a-d, g-j, l, n-p**) and bars indicate mean ± SEM. Statistical differences are depicted as *p < 0.05, **p < 0.01, ***p < 0.001.

Mast cells are known to expand upon helminth infection and can be found in the skin and other barrier tissues. ^12, 14^ We detected a clear expansion of mast cells in the spleen after *H. polygyrus* infection (Supplementary Fig. 2b-2d), but no difference was found in the skin (Fig. 2g, Supplementary Fig. 2e, 2f). Innate lymphoid cells (ILCs) and γδ T cells are common in barrier tissues in mice, but there was no expansion of either cell type in the skin after worm infection (Fig. 2h, 2i, Supplementary Fig. 2g, 2h).

Mouse skin comes into contact with faeces and intestinal helminths have been shown to increase intestinal permeability and bacterial translocation from the gut. ^2^ We next tested if T cell priming by intestinal bacteria in combination with environmental skin exposure to bacteria of faecal origin could drive T cell accumulation in skin. Indeed, *H. polygyrus-*infected mice had elevated levels of soluble CD14 (sCD14) in serum (Supplementary Fig. 2i), an indirect marker for increased levels of LPS in blood, ^15^ suggesting enhanced bacterial translocation from the gut to the blood. To investigate if the increase in CD4+ cells in skin was due to bacterial translocation, we measured CD4+ cell numbers in the skin of mice with colitis induced by dextran-sodium sulphate (DSS) treatment, which is known to increase LPS levels in the blood. ^16^ DSS-induced colitis was established over a timeline similar to chronic *H. polygyrus* infection (Supplementary Fig. 2j). This protocol did not increase skin CD4+ cell numbers (Fig. 2j), indicating that bacterial translocation and generalized inflammation was not the cause of higher CD4+ numbers in the skin.

To dismiss that worm products excreted in mouse faeces caused T cell recruitment to skin, *H. polygyrus*-infected mice were dewormed early after infection when adult worms have left the intestinal wall but not started to produce eggs. ^12^ Cages were changed after deworming to minimize skin exposure to faecal antigens. Dewormed mice were then co-housed with heavily *H. polygyrus-*infected mice or uninfected controls (Fig. 2k). No difference in CD4+ T cell numbers was detected in skin collected from these groups (Fig. 2l; HpDw and HpDw CoH). However, in line with previous experiments, CD4+ T cell numbers were increased in previously infected mice compared to uninfected controls irrespective of co-housing partners (Fig. 2l). This indicates that the gut infection rather than environmental worm antigen is responsible for CD4+ T cell accumulation in the skin.

### Skin-homing receptors are up-regulated on circulating CD4+ T cells after H. polygyrus infection

Tissue tropism is induced during priming of T cells by expression of homing receptors. ^17^ T cell trafficking to the skin depends on the expression of cutaneous lymphocyte-associated antigen (CLA) and the chemokine receptors (CCR) CCR4 and CCR10. ^17^ As expected, the mesLNs showed a great expansion of CD4+ cells after *H. polygyrus* infection (Supplementary Fig. 3a-3c). Interestingly, expression of skin-homing receptors CCR4 and CCR10 was upregulated in CD4+ cells from the mesLNs of worm-infected mice 2 weeks after infection (Fig. 3a-3h, Supplementary Fig. 3d-3f). CCR4+ and CCR10+ CD4+ T cells were also increased in blood after infection (Fig. 3i-3n, Supplementary Fig. 3g). CCR9+ cells were also upregulated in the blood (Supplementary Fig. 3h-3i), in line with CCR9 being induced during gut infection to target cells to the intestine. ^17^ This indicates that CD4+ T cells activated by *H. polygyrus* in mesLN are directed to extraintestinal tissues such as the skin through upregulation of skin-homing receptors.

**Figure 3:**
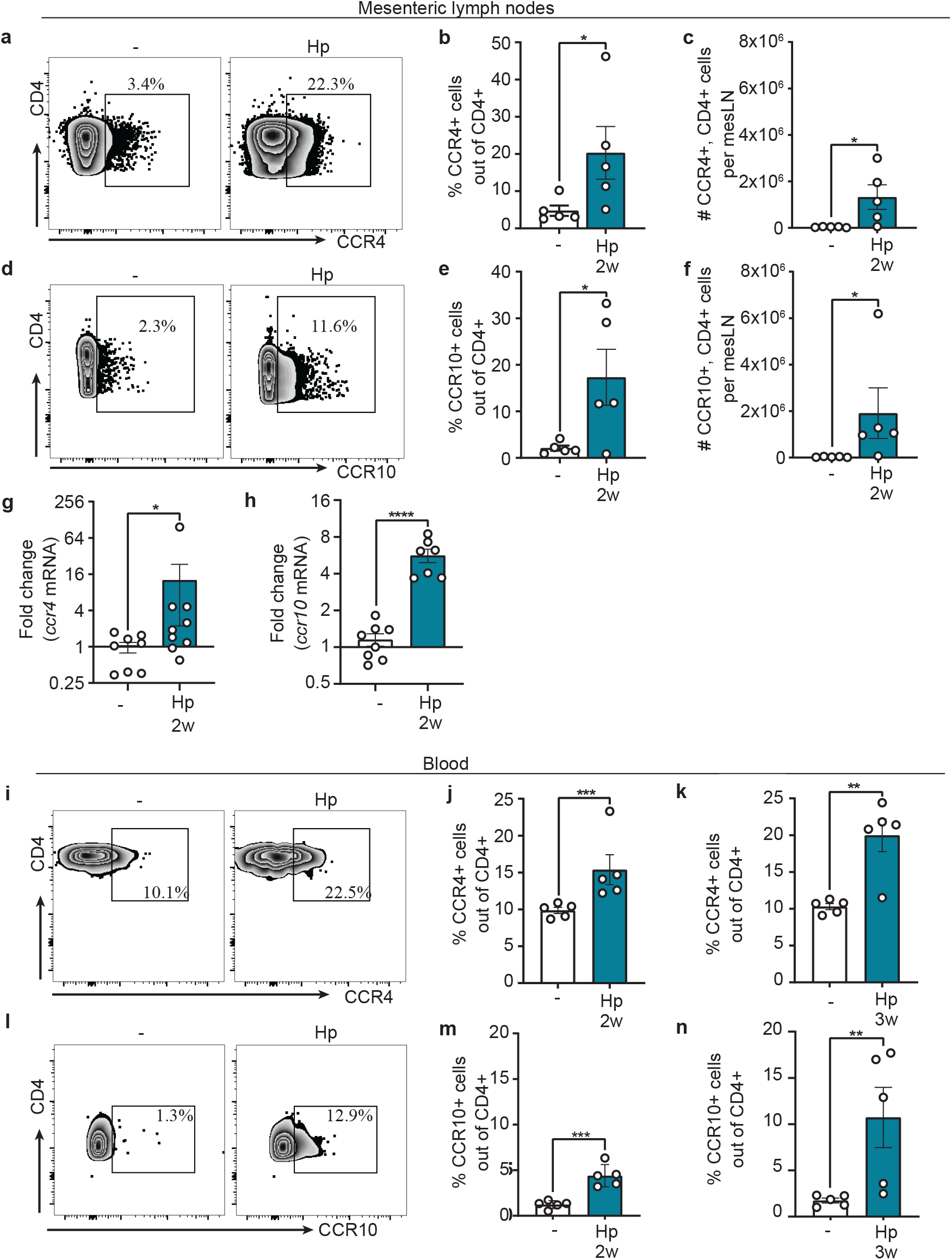
Skin-homing receptors are up-regulated on mesLN and circulating CD4+ T cells after *H. polygyrus* infection. **a-n** Mice were infected with *H. polygyrus* (Hp) or not (-). **a-h** Mesenteric lymph nodes (mesLNs) collected two weeks after infection. **a** Representative FACS plots of CCR4 staining gated on CD4+ mesLN cells. **b, c** Frequencies and absolute numbers of CCR4+ cells out of CD4+ cells. **d** Representative FACS plots of CCR10 staining gated on CD4+ mesLN cells. **e, f** Frequencies and absolute numbers of CCR10+ cells gated on CD4+ cells. **g, h** Fold change of CCR4 and CCR10 mRNA expression in CD4+ T cells bead isolated from mesLNs. **i-n** Blood collected at indicated time points after infection. **i** Representative FACS plots of CCR4 staining gated on CD4+ blood cells. **j, k** Frequencies of CCR4+ cells out of CD4+ cells. **l** Representative FACS plots of CCR10 staining gated on CD4+ cells. **e, f** Frequencies and absolute numbers of CCR10+ cells out gated on CD4+ blood cells. Samples were analysed by flow cytometry (**a-f, i-n**) or qPCR (**g-h**). In **b, c, e, f, j, k, m, n**, one out of at least two independent experiments with similar results are shown. **g, h** shows pooled data from two independent experiments. Each dot represents one mouse (n≥5) and bars indicate mean ± SEM. Statistical differences depicted as *p < 0.05, **p < 0.01, ***p < 0.001, **p < 0.01, ****p < 0.0001.

### T_H_2 cells accumulate and persist in the skin after H. polygyrus infection

Isolated studies have demonstrated that T_H_2 cells expand during worm infection in the gut and also accumulate at other sites where they persist for at least 2 weeks after de-worming. ^18, 19^ Next, we characterized the phenotype of CD4+ T cells infiltrating the skin after *H. polygyrus* infection. The frequency of T_H_2 cells (ST2+, Foxp3-CD4+ ^20^) was clearly increased in the skin both at 2 and 4 weeks after infection (Fig. 4a, 4b, Supplementary Fig. 4a), while no increase in T_REG_ cells (Foxp3+ CD4+) was detected at either time point (Fig. 4c, Supplementary Fig. 4b, 4c). Accordingly, skin of worm-infected mice contained a higher frequency of CD4+ cells that produced the T_H_2-associated cytokines IL-4 and IL-13 upon polyclonal re-stimulation (Fig. 4d, 4e). Whereas CD4+ T cells were clearly skewed towards a T_H_2 phenotype, there was no skewing of ILCs towards an ILC2 phenotype (Fig. 4f, Supplementary Fig. 4d). In the blood, T_H_2 cells were expanded 2 weeks after infection, but had started to contract 2 weeks later (Fig. 4g). Interestingly, out of CD4+ blood T cells, ST2+ cells expressed higher levels of CCR4 and CCR10 skin-homing chemokine receptors compared to ST2-cells, a difference that was more pronounced in *H. polygyrus*-infected mice (Fig. 4h, 4i). T_H_2 cells seem therefore to be especially targeted to the skin in worm-infected mice.

**Figure 4:**
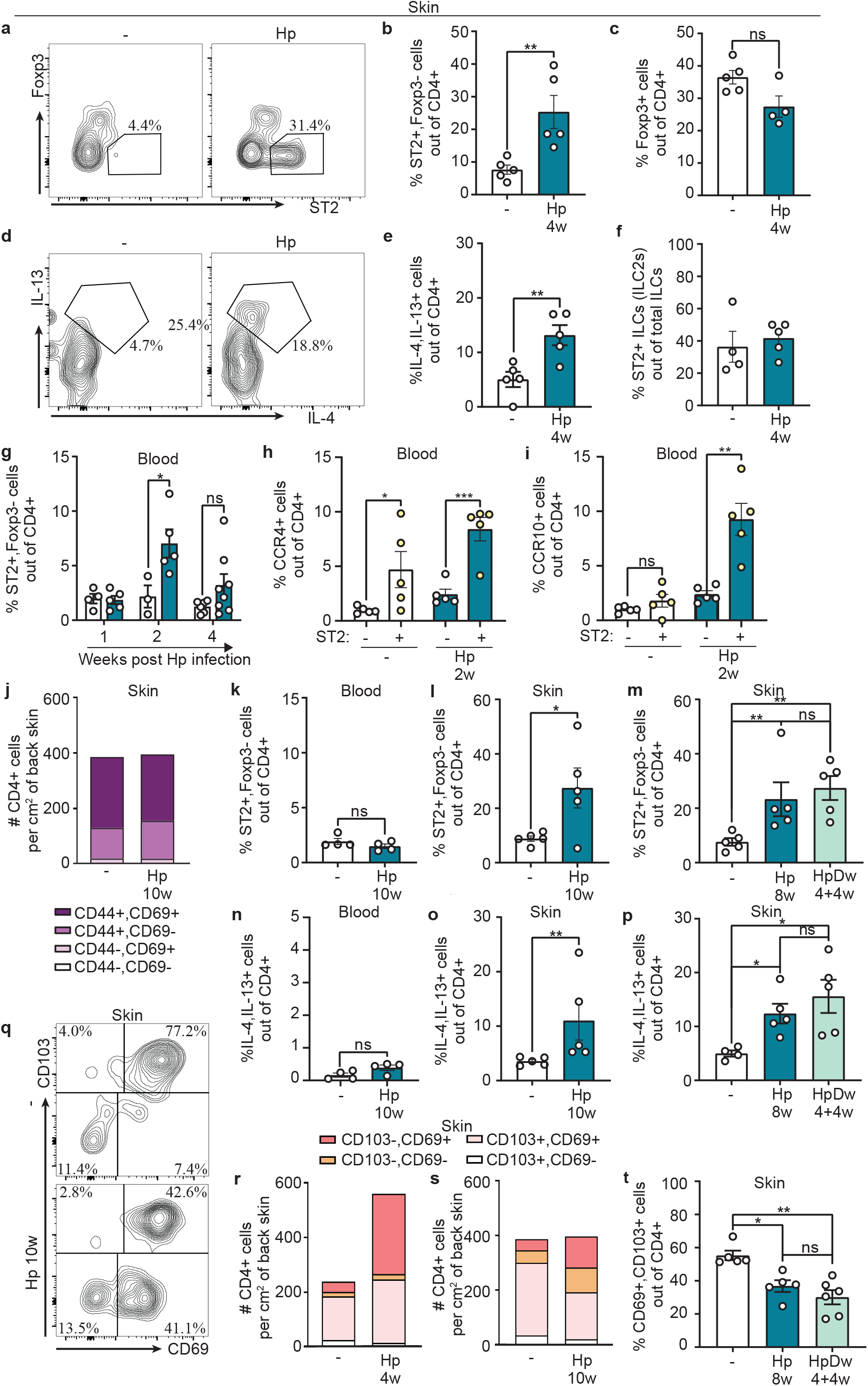
Type 2 CD4+ T cells accumulate and persist in skin after *H. polygyrus* infection. **a-t** Mice were infected with *H. polygyrus* (Hp) or not (-). a-f Flow cytometry on back skin. **a** Representative FACS plots of ST2+, Foxp3-staining gated on CD4+ cells. **b** Frequency of ST2+, Foxp3- and **c** Foxp3+ cells out of CD4+ cells. **d** Representative FACS plots of IL-13+, IL-4+ cells gated on CD4+ T cells. **e** Frequency of IL-13+, IL-4+ cells out of CD4+ cells after PMA/ionomycin restimulation in the presence of Brefeldin A. **f** Frequency of ST2+ cells out of ILCs. **g-i** Flow cytometry on blood. **g** Frequency of ST2+, Foxp3-cells out of CD4+ T cells. **h** Frequency of CCR4+ and i CCR10+ in the ST2 expressing (+) and non-expressing fraction (-) of CD4+ T cells. **j** Absolute numbers and percentages of CD44 and CD69 out of CD4+ T cells. Data is calculated from an average of three independent experiments with a total of ≥12 mice per group. **k-m** Frequencies of ST2+, Foxp3-cells out of CD4+ cells in **k** blood and **l, m** skin, in **m** in mice dewormed 4 weeks before analysis. **n-p** Frequencies of IL-13+, IL-4+ cells out of CD4+ cells in **n** blood and **o, p** skin, in **p** in mice dewormed 4 weeks before analysis. **q** Representative FACS plots of CD103 and CD69 gating in CD4+ T cells. **r, s** Absolute numbers and percentages of CD103 and CD69 out of CD4+ T cells. Data is calculated from an average of three independent experiments with a total of ≥12 mice per group. **t** Frequency of CD103+, CD69+ cells out of CD4+ T cells, mice were dewormed 4 weeks before analysis. **a-t** One out of at least two independent experiments with similar results are shown. Each dot represents one mouse (n≥4, **b-c, e-i, k-p, t**) and bars indicate mean ± SEM. Statistical differences are depicted as *p < 0.05, **p < 0.01, ***p < 0.001.

To address if worm-induced accumulation of CD4+ T cells in the skin persisted over time and following deworming treatment, we assessed skin CD4+ T cells for up to 10 weeks after infection, when most of our animals have naturally cleared infection (Supplementary Fig. 4e). At this timepoint the number of T cells had normalized in the skin and the frequency of activated cells (CD44+/CD69+) was similar to those of uninfected and more recently infected mice (Fig. 4j, 2f). Although there were no signs of elevated circulating T_H_2 cells in worm-infected mice, a higher frequency of T_H_2 cells persisted in the skin of worm-infected animals (Fig. 4k, 4l, 4n, 4o). Likewise, animals from which worms had been removed by drug treatment 4 weeks earlier retained higher frequencies of T_H_2 cells in the skin compared to uninfected animals. The frequency of T_H_2 cells in dewormed mice did not differ from that of animals still carrying the infection (Fig. 4m, 4p, Supplementary Fig. 4e).

Due to the longevity of T_H_2 cells in the skin we next assessed the expression of memory markers on skin CD4+ T cells in our model. CD103 and CD69 are established T_RM_ markers for CD8+ T cells but no reliant markers have yet been established for tissue residency of CD4+ T cell and occurrence of recirculation is debated. ^7, 8, 11^ We found that the CD103-, CD69+ population of CD4+ T cells expanded early after *H. polygyrus* infection (Fig. 4r). Surprisingly, this subset persisted when total cell numbers normalized 10 weeks post infection (Fig. 4q, 4s) and 4 weeks after deworming, leaving fewer CD69+CD103+ cells in the skin of long-term infected- and dewormed-mice (Fig. 4q, 4s, 4t). CCR8 have recently been suggested to define a CD4+ T_RM_ cell population in human skin, ^21^ but CCR8 was also reduced on skin CD4+ cells late after *H. polygyrus* infection or deworming (Supplementary Fig. 4f). Moreover, we did not find changes in CD11a and KLRG-1 (Supplementary Fig. 4g, 4h), which are associated with long-lived CD4+ cells. ^22^ In summary, our data show that activated T_H_2 cells accumulate and persist in the skin of mice infected with *H. polygyrus* but lack several markers associated with T_RM_ cells.

### Skin CD4+ cells are H. polygyrus-specific

To investigate the reactivity of skin-localized CD4+ T cells in *H. polygyrus-*infected mice, we injected soluble worm antigen (SWAg) from *H. polygyrus* into the back skin of infected and uninfected mice and analysed cytokine mRNA expression by qPCR 24 hours later. Transcripts for IL-4, IL-5 and IL-13 were clearly induced by SWAg in *H. polygyrus-*infected mice whereas no induction was seen in uninfected controls (Fig. 5a-5c). We did not detect induction of T_REG_ or T_H_1-associated cytokines after SWAg injection (Supplementary Fig. 5a-5d). To assess direct recall in the skin we measured footpad swelling elicited by SWAg. Indeed, worm-infected animals displayed a swelling reaction upon SWAg administration in the footpad skin, which was absent in uninfected mice (Fig. 5d). This reaction was MHC-II-dependent (Fig. 5d) and hence mediated by CD4+ T cells. We then depleted circulating lymphocytes using FTY720 which prevents lymphocyte egress from LNs and tissues. ^6^ FTY720 treatment depleted lymphocytes from the circulation (Supplementary Fig. 5e) but did not affect SWAg-induced swelling in worm-infected mice (Fig. 5e), indicating that persisting local skin CD4+ T cells mediated the reaction. Many intestinal helminths infect by penetrating the skin, including the murine hookworm *Nippostrongylus brasiliensis*. ^23^ *H. polygyrus* infection has previously been shown to protect against subsequent challenge with *N. brasiliensis* at the skin-penetration stage. ^23^ We next injected antigen from *N. brasiliensis* in the footpad and measured footpad swelling, but no swelling reaction was observed in *H. polygyrus*-infected mice (Fig. 5f, 5g), highlighting antigen specificity of the accumulated CD4+ T cells. In summary, CD4+ cells that accumulate in the skin of infected mice are *H. polygyrus*-specific and respond to *H. polygyrus* in an MHC-II-dependent manner by producing T_H_2-associated cytokines.

**Figure 5:**
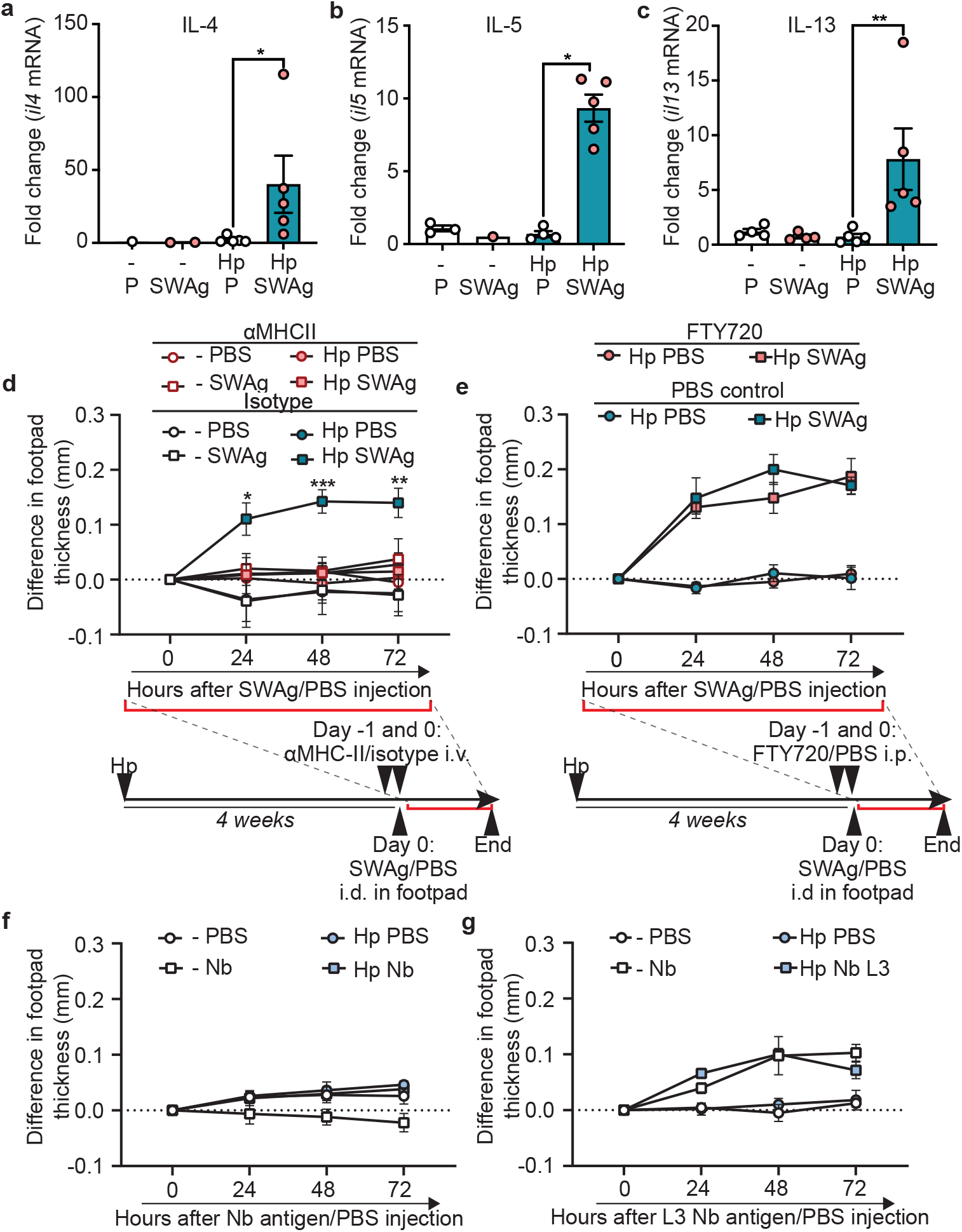
Skin CD4+ cells display antigen-specific responses to *H. polygyrus*. Mice were infected with *H. polygyrus* (Hp) or not (-). **a-c** Soluble worm antigen from (SWAg) *H. polygyrus* (and PBS (P) in contralateral side as control) was injected in the back skin four weeks p.i. Fold change of mRNA expression of indicated cytokines analysed by qPCR. 5 mice per group were always used, and when fewer dots are shown expression was under detection levels. **d** Footpad swelling after SWAg injection in anti-MHC-II antibody-treated and untreated mice and in **e** FYT720-treated and untreated mice. **f** Footpad swelling after injection of antigen from adult *N. brasiliensis* (Nb) or **g** stage L3 *N. brasiliensis* larvae (L3 Nb), 4 weeks pi. PBS was injected in the contralateral side as control. **a-g** One out of at least two independent experiments with similar results are shown. Each dot represents a mouse (**a-c**) or the mean ± SEM (**d-g**) of a group of at least five mice and statistical differences depicted are as *p < 0.05, **p < 0.01, ***p < 0.001. In d and e, comparisons are made to the Hp-infected, SWAg injected but not treated with αMHCII (isotype, **d**) or FTY720 (**e**), respectively.

### H. polygyrus-specific footpad swelling and impaired IFN-γ responses to mycobacterial antigens remain long after deworming

Our previous studies showed that deworming can restore immune reactivity of the skin-draining LNs dampened by *H. polygyrus* infection. ^6^ To test if the altered composition of T cells seen weeks after clearance of worms would affect subsequent responses, we injected SWAg in the footpad 12 weeks after *H. polygyrus* infection or 10 weeks after deworming (Fig. 6a). Mice that had been infected with *H. polygyrus* retained a noticeable skin reaction upon SWAg injection at both time points (Fig 6a). Furthermore, in mice previously infected but dewormed 10 weeks prior to WCL injection in the skin, IFN-*γ* recall responses were still impaired at both WCL-injected and untouched skin sites (Fig. 6b-6d). This implies that *H. polygyrus* infection in the gut causes durable changes in the composition of skin T cells and persistently alters the capacity to form skin responses to an unrelated antigen (Fig. 6e).

**Figure 6:**
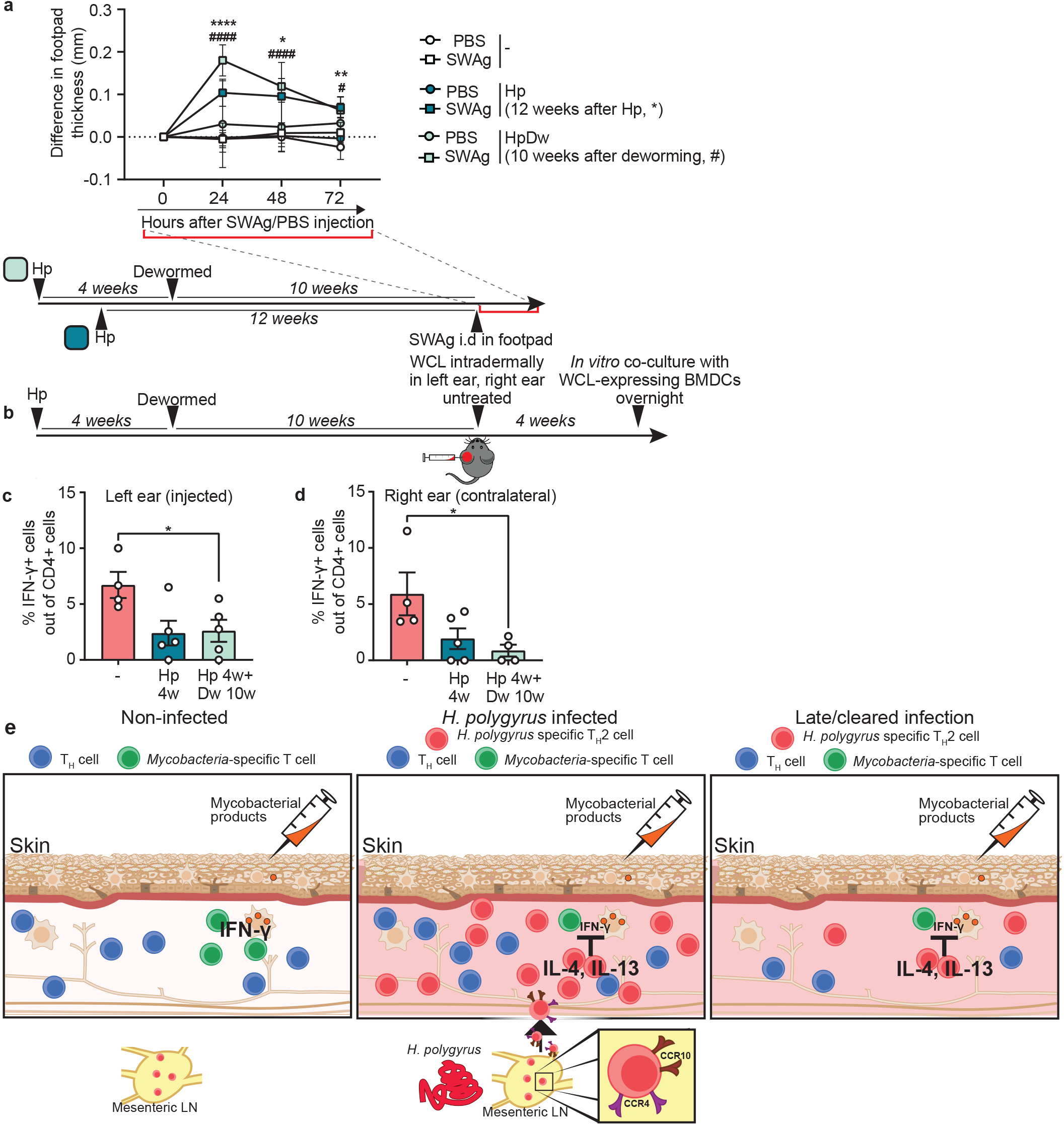
*H. polygyrus*-specific footpad swelling and muted skin IFN-γ responses to mycobacterial antigens remain long after removal of worms. Mice were infected with *H. polygyrus* (Hp) or not (-). **a** Footpad swelling after soluble worm antigen (SWAg) from *H. polygyrus* injection in the footpad 12 weeks p.i. or 10 weeks after deworming. **b** shows the setup for **c** and **d**: Mice were dewormed four weeks after Hp infection and 10 weeks later whole cell lysate (WCL) from *M. tuberculosis* was injected in the left ear and the right ear was left untreated. Four weeks later, ears were collected, and ear cells cultured with BMDCs expressing WCL overnight and analysed by flow cytometry. **c-d** Frequency of IFN-γ+ cells out of CD4+ T cells. **a-d** One out of at least two independent experiments with similar results are shown. Each dot represents the mean ± SEM of a group of mice of n≥4, (**a**) or an individual mouse (**c, d**) and bars indicate mean ± SEM. Statistical differences are depicted as *p < 0.05, **p < 0.01, ****p < 0.0001 or in a as above for 12 weeks Hp infected mice and as #p < 0.05, ####p < 0.0001 for mice dewormed 10 weeks prior to antigen injection. **e** shows a graphical summary model of our findings: When worms are not present in the gut, mycobacteria-reactive CD4+ T cells accumulate in the skin after WCL injection and are able to respond appropriately with IFN-**γ** production to re-stimulation by mycobacterial products. Intestinal Hp infection causes up-regulation of skin-homing receptors on CD4+ T cells, and skin infiltration of *H. polygyrus*-specific T_H_2 cells from intestine that dampen IFN-γ production in response to mycobacterial products. These T_H_2 cells remain in the skin after the intestinal infection is cleared and total cell numbers have normalized, persistently transforming the skin CD4+ T cell pool.

## Discussion

Intestinal worm infection impact immune responses to other pathogens, vaccines, autoantigens, allergens and contact sensitisers. ^2, 6, 13, 24^ These effects have been believed to be due to expansion of T_REG_ ^2, 24, 25^ and/or T_H_2 cells, ^18, 26^ atrophy of peripheral LNs ^6, 27^ or changes in the intestinal microbiota or its metabolome. ^28-30^ Yet, the full spectrum of helminth-induced effects on the immune system is not elucidated. In our study we show that infection with the intestinal nematode *H. polygyrus* promote homing to and long-term residence of *H. polygyrus*-specific T_H_2 cells producing IL-4 and IL-13 in the skin. We propose that this gut infection-induced remodelling of the skin T cell landscape led to dampened skin recall responses to unrelated antigens, both locally and distally to the injection site. The impaired skin T cell response persisted after *H. polygyrus* had been cleared from the host. IFN-*γ* responses in other organs have previously been shown to be muted in helminth-infected humans and animals. ^4, 13, 31^ For example and in line with our findings, intestinal helminth-induced suppression of immunity towards *M. tuberculosis* has been shown to be mediated via the common IL-4/IL-13 receptor (IL-4Rα) pathway in the lung. ^26^ Our findings add to the growing body of literature describing how worms can form our immune system and cause extraintestinal immunosuppression ^2, 6, 13, 18, 24, 30^ and propose new ways by which gut-restricted worms impact the onset of skin immune responses.

A number of mechanisms have been suggested for cell trafficking into tissues distinct from the infection site, many focusing on mucosal tissues. ^32-37^ Studies have proposed the dissemination of pathogen products, ^32^ re-programming of lymphocyte homing profile by a new tissue or LN microenvironment ^33-36^ and upregulation of chemokine receptors on cells in LNs draining the infection site that target them to sites distal to that of the infection. ^37^ Recruitment of CD4+ T cells into skin was shown not to be a consequence of bacterial translocation into the blood or excretion of worm antigens into the environment. Instead, *H. polygyrus* infection triggered expression of skin-homing CCR4 and CCR10 on CD4+ T cells, indicating a more direct link between gut and skin. Our data indicate that the higher number of T_H_2 cells lodged in the skin of *H. polygyrus*-infected animals is a result of induction of skin-homing chemokine receptor expression on CD4+ T cells in the mesLN followed by travelling through the circulation.

The functional advantage of the observed recruitment of CD4+ T cells into skin is not clear. Since a number of worm species infect through the skin, ^38^ skin-homing T_H_2 cells could provide direct protection against repeated exposure to larval forms of the same skin-infecting nematode or cross-protection against other worms. Indeed, *H. polygyrus* can protect against challenge by the *N. brasiliensis* ^23, 39^ and cross-protection between helminth species have also been seen in other models. ^40, 41^ However, we did not observe cross-reactivity to *N. brasiliensis* antigens in our model as measured by footpad swelling. Prior studies point to innate type 2 responses rather than CD4+ T cells as mediators of such cross-reactivity. ^23, 40, 41^ We did not find any differences in ILCs, which might explain the lack of cross-reactivity. Our skin responses were MHC-II-dependent, did not require circulation of lymphocytes and produced IL-4 and IL-13 production after *in vitro* restimulation with SWAg, revealing a strong *H. polygyrus*-specific T_H_2 component among the CD4+ T cells accumulating in the skin. Importantly, T_H_2 cells persisted in skin even after the expulsion of *H. polygyrus* from the gut and skin recall responses to a secondary challenge with a mycobacterial antigen preparation were muted long after worm expulsion. The changed composition of skin CD4 T cells with compromised ability of the skin of worm-infected mice to mount robust reactions to subsequent antigen challenge have consequences for immune responses to pathogens and vaccines. Intriguingly, our results provide a novel explanation to the geographically mutually exclusive high prevalence of inflammatory skin disorders and the responsiveness to the intradermally administered tuberculosis vaccine BCG, observed around the world. ^3, 4, 31^

## Methods

### Mice and ethics

Three to 4 weeks old C57Bl/6NRj mice were obtained from Janvier Labs (France), Taconic Biosciences (Denmark) or Karolinska Institutet and acclimatized for at least 1 week before start of experiment. Mice were housed and handled under SPF conditions at the BSL-2 facility at Comparative Medicine Biomedicum (KM-B) according to Swedish national regulations for laboratory animal work with free access to food and water, cage enrichment (nesting material, gnaw stick) and 12 hours light and dark cycles. Mice were anaesthetised with isoflurane during skin injections. Female mice were used in most experiments. Experimental group size was determined based on previous experience of variation (often one cage of 5 animals, as provided to us) and mice/cages allocated randomly into groups. Experiments were not blinded at any stage. Experiments were granted by the regional ethical board (Stockholms djurförsöksetiska nämnd) under permit numbers 6738/19, N89/15 and N171/14 with the amendment N131/16 and exemption from L150 Dnr 5.2.18-7344/14 and approved by the Swedish Board of Agriculture.

### Skin collection

Back skin of euthanized mice was shaved and Veet® hair removal cream was applied onto the shaved area. Cream was removed after 3-5 minutes and the treated area washed with PBS. Back skin was collected with biopsy punches. Ears were removed with scissors without prior shaving or treatment.

### Infections and in vivo recall responses

Mice were infected with 200 or 300 L3 stage larvae of *H. polygyrus* per oral gavage at 4 to 5 weeks of age, unless otherwise stated in figure legends. At the end of each experiment, small intestines were collected, opened longitudinally, placed in a mesh in RPMI-1640 and incubated at 37°C for 2-4 hours. Live worms migrated out and were collected from the bottom of the tube for counting and antigen preparations (see below). Deworming was done with 2 mg Fyrantel® paste (0.8 mg pyrantel pamoat) per mouse for 3 constitutive days and successful treatment was confirmed by lack of eggs in the faeces. Whole cell lysate (WCL) from *M. tuberculosis* strain H37Rv was obtained from BEI Resourses, NIAID, NIH. Ears were injected intradermally with 5 µg WCL in 5 µl PBS. For recall responses, worm antigen was injected in footpads and footpad swelling measured with a digital caliper or in shaved and Veet®-treated back skin for cytokine assessment by qPCR. Worm antigen was administered in 20 µl in back skin or footpad at 50 µg (SWAg or adult *N. brasiliensis*) or 25 µg (L3 *N. brasiliensis*).

### Worm antigens

Soluble worm antigen (SWAg) from *H. polygyrus* and antigens from *N. brasiliensis* was prepared as previously described ^13, 42^ and stored at -80°C.

### Dextran sodium sulphate model

Dextran sodium sulphate (DSS, TdB Consultancy AB) was provided at 2% w/v *ad libitum* in the drinking water for 2 periods of 7 days each with a recovery period of 14 days in between. Mice were sacrificed 7 days following the second DSS period.

### Antibody and FTY720 treatments

Anti-MHC-II was injected intravenously and Fingolimod/FTY720 (3 mg/kg, Sigma Aldrich) intraperitoneally the day before and just before SWAg footpad injection.

### CD4+ T cell bead isolation

MesLNs were collected and CD4+ T cells isolated using the Dynabeads™ Untouched™ Mouse CD4 Cells Kit (Thermo Fisher) according to the manufacturer’s instructions.

### Gene expression by qPCR

Four mm back skin biopsies were collected into QIAzol™ lysis reagent (Qiagen) and chopped into smaller pieces. Skin samples were homogenized in a TissueLyser LT (Qiagen, 50 Hz, 2 times 4 minutes) using 5 mm stainless steel beads (Qiagen). RNA was extracted by the chloroform/isopropanol method and converted to cDNA as previously described. ^13^ qPCR was performed using TaqMan-technology (Applied Biosystems) with the best matched primers and probe kits from Life technologies and run on a CFX384 qPCR machine (BioRad). Target gene expression data was normalized to expression of β-actin for individual samples and fold change over control samples calculated by using the 2^-ΔΔCt^ method.

### Flow cytometry

Back skin and ears were cut into smaller pieces and incubated in collagenase 3 (3 mg/ml, Worthington), DNAse I (5 µg/ml, Roche) and 10% FBS (Sigma) in RPMI-1640 at 37°C, 5% CO_2_ for 90 minutes. Samples were then grinded in BD Medimachine for 4 minutes, filtered, and cells washed in PBS. Blood was collected into EDTA-containing tubes (Sarstedt), red blood cells (RBCs) lysed and cells washed in PBS. Spleens were homogenized through 70 µm strainers with syringe plunger, RBCs lysed, and cells washed in PBS. Single cell suspensions were stained with LIVE/DEAD™ Fixable Yellow Dead Cell Stain according to manufacturers’ instructions (Invitrogen), washed in FACS buffer (4% FBS, 3 mM EDTA, and 40 μg/ml DNAse I (Roche)) and incubated with combinations of antibodies to CD45 (30-F11), CD4 (RM4-5 or GK1.5), CD8 (53-6.7), KLRG-1 (2F1), CD44 (IM7) from BD, TCRβ (H57-597), PD-1 (29F.1A12), ST2 (DIH9), CCR8 (SA214G2), CCR4 (2G12), CD69 (H1.2F3), CD103 (2E7), γδ TCR (GL3) from Biolegend, CCR10 (248918) from R&D, CD11a (M17/4) from eBioscience, FcεRI (MAR-1), c-Kit (CD117, 2B8) from Invitrogen and Mouse BD Fc Block™, for 30 minutes at 4°C. To assess T_H_2 cytokine production, PMA (50 ng/ml, Sigma), ionomycin (5 μM, Sigma) and Brefeldin A (5 μg/ml, Sigma) was added to the cells for 3 hours. For WCL restimulation, bone marrow-derived dendritic cells were collected as previously described, ^43^ incubated with WCL (10 mg/ml) overnight, and ear cells were added the following day. The co-culture was incubated with WCL overnight with Brefeldin A added for the last 3 hours. Stimulated cells were fixed and permeabilized with the Foxp3 staining kit (eBioscience) or IntraPrep Permeabilizaton Reagent (Beckman Coulter) and then stained with combinations of Foxp3 (FJK-16s) and IL-13 (eBio13A) from Invitrogen and IL-4 (11B11) and IFN-γ (XMG1.2) from BD. Cells were analysed on LSR II flow cytometer (BD). Absolute numbers of cells were calculated by using CountBright™ Absolute Counting Beads (Invitrogen) or manual counting in haemocytometer and data analysed in FlowJo V10 (BD).

### Soluble CD14 ELISA

CD14 levels were determined in mouse serum by ELISA kit (Biometic Gmbh) according to the manufacturer’s instructions.

### Statistics and data presentation

Outliers plausibly appearing from technical errors or unknown reasons were excluded prior to analysis by the ROUT test as recommended by GraphPad Prism to eliminate subjectivity when evaluating validity of data, as determined *a priori*, as confounders were not explicitly controlled. For statistical comparison between 2 groups, unpaired student’s *t* test was performed when data followed normal distribution and Mann-Whitney U when distributions did not. One-way ANOVA with Tukey post-test was used for multiple comparisons. GraphPad Prism version 8 or 9 were used.

## Acknowledgement

We thank Juan Basile and Cassandra Hokka-Zakrisson for technical assistance. Flow cytometry was performed at the Biomedicum Flow Cytometry Core facility (BFC), Department of Microbiology, Tumor and Cell Biology (MTC), Karolinska Institutet. This work was supported by Karolinska Institutet and Vetenskapsrådet (VR). ER was funded by the Wenner-Gren Foundations. The funders had no role in study design, data collection and interpretation or the decision to submit the work for publication.

## Author contributions

SN and CC conceived and designed the project. ML, JM, ALC, XF, CT, JS and RC performed experiments and CC analysed the data. ER, AGR, LB and EV provided essential tools. AGR, EV, LE, JC provided advice on experimental design and conceptualization. CC, AGR and SN wrote the paper and all authors reviewed and approved the final version of this manuscript to be published.

