## Supplementary material for "Intestinal helminth infection transforms the CD4+ T cell composition of the skin"

### Supplementary figure 1

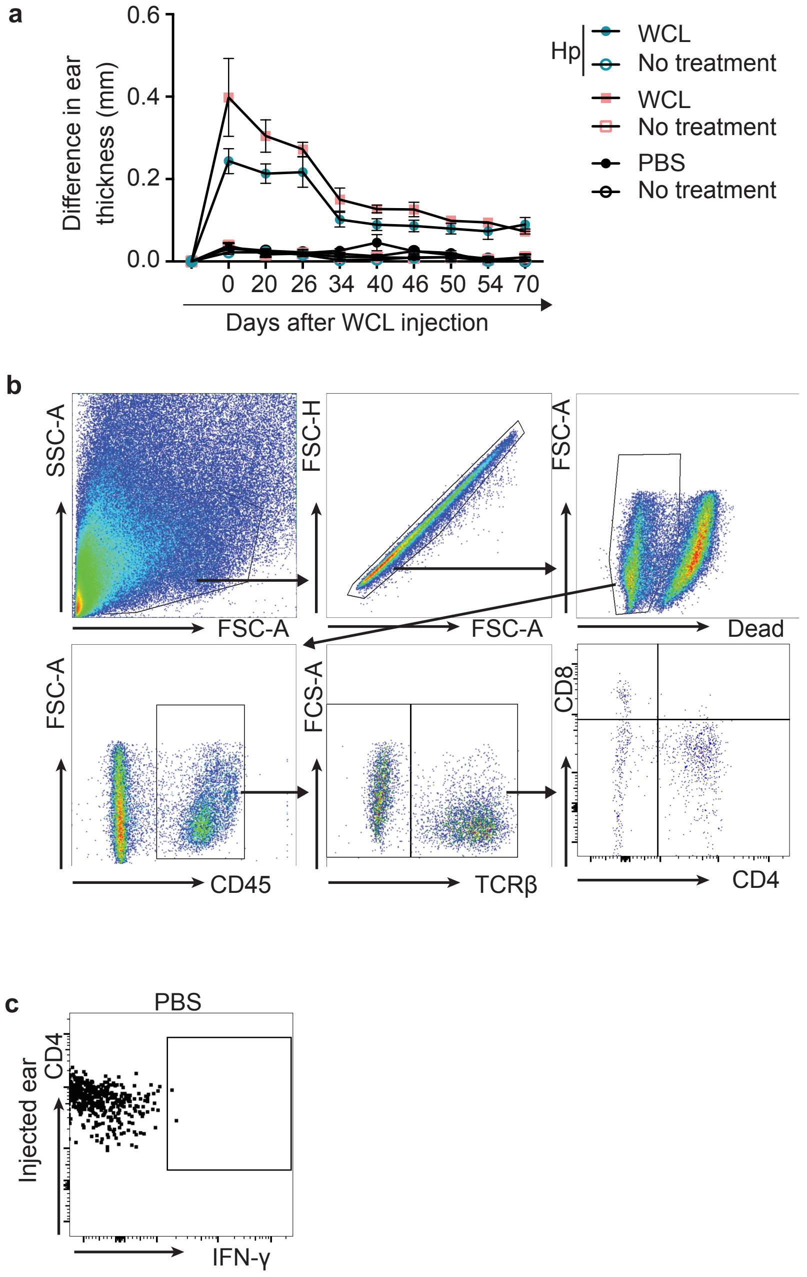


**Supplementary Figure 1: Complement to Figure 1. a** Ear thickness measured after WCL injection in ears of non-infected and *H. polygyrus*-infected (Hp) mice. Each dot represents the mean ± SEM of a group of five mice. **b** shows example of gating used for CD4+ T cells in ears. In **c**, an example of a PBS injected ear is shown as gate reference for data in Fig. 1b-1h.

### Supplementary figure 2

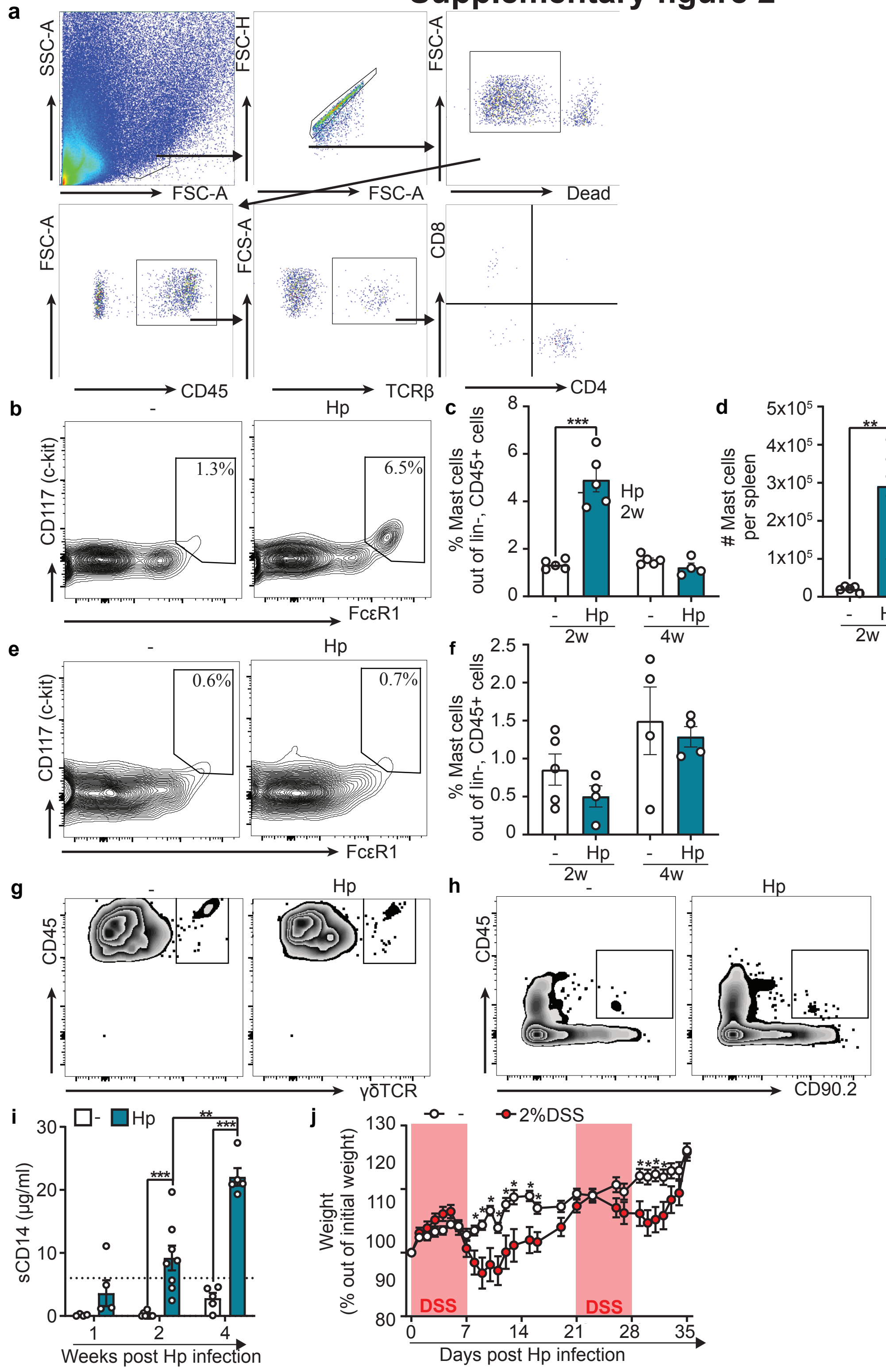


**Supplementary Figure 2: Complement to Figure 2. a** Example of gating used for CD4+ T cells in back skin. **b-i** Mice were infected with *H. polygyrus* (Hp) or not (-). **b** Representative FACS plot of mast cells (c-Kit+, FcεRI+, out of linage- (lin-), CD45+ cells) in spleen. **c, d** Frequencies and absolute numbers of mast cells in spleen. **e** Representative FACS plot of mast cells in back skin. **f** Frequencies of mast cells in back skin. **g** Representative FACS plot of γδTCR+ T cells in back skin. **h** Representative FACS plot of innate lymphoid cells (ILCs, lin-, CD45+, CD90.2+) in back skin. **i** Soluble (s) CD14 levels measured by ELISA in serum. The dotted line represents the normal cut-off value for an uninfected mouse. **j** Weight of mice treated with 2% dextran sodium sulfate (DSS) in the drinking water and indicated time points after start of treatment. Periods of treatment are shown in gray. **a-i** One out of at least two independent experiments with similar results are shown. In **j**, the experiment has been performed once but with 10 mice (divided in 2 cages) per group. Each dot represents an individual mouse (**c-d, f, i**) or the mean ± SEM of a group of mice of (n=10, **j**) and bars indicate mean ± SEM. Statistical differences are depicted as *p < 0.05, **p < 0.01, ***p < 0.001, ****p < 0.0001.

### Supplementary figure 3

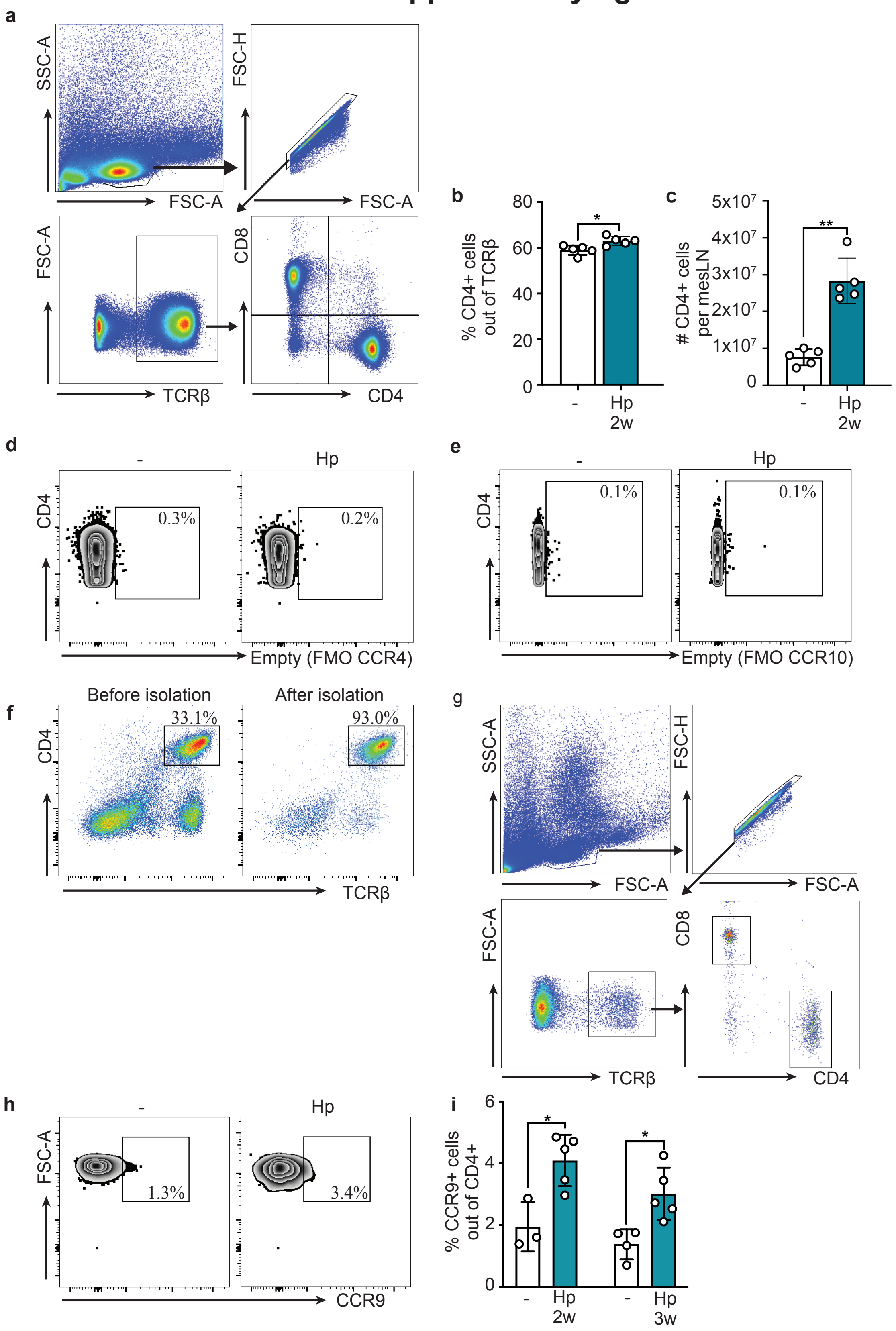


**Supplementary Figure 3: Complement to Figure 3.** Mice were infected with *H. polygyrus* (Hp) or not (-). **a** Example of gating used for CD4+ T cells in mesenteric lymph nodes (mesLNs). **b, c** Frequency and absolute number of CD4+ T cells in mesLNs. **d, e** FMO controls for CCR4 and CCR10 staining, respectively. **f** shows the purity of CD4+ T cells after bead isolation for qPCR analysis in Fig. 3g and 3h. **g** Example of gating used for CD4+ T cells in blood. **h** Representative FACS plot of CCR9+ staining in blood. **i** Frequencies of CCR9+ cells out of blood CD4+ T cells. One out of at least two independent experiments with similar results are shown. Each dot represents a mouse (n≥3, **b-c, i**) and bars indicate mean ± SEM. Statistical differences are depicted as *p < 0.05, **p < 0.01.

### Supplementary figure 4

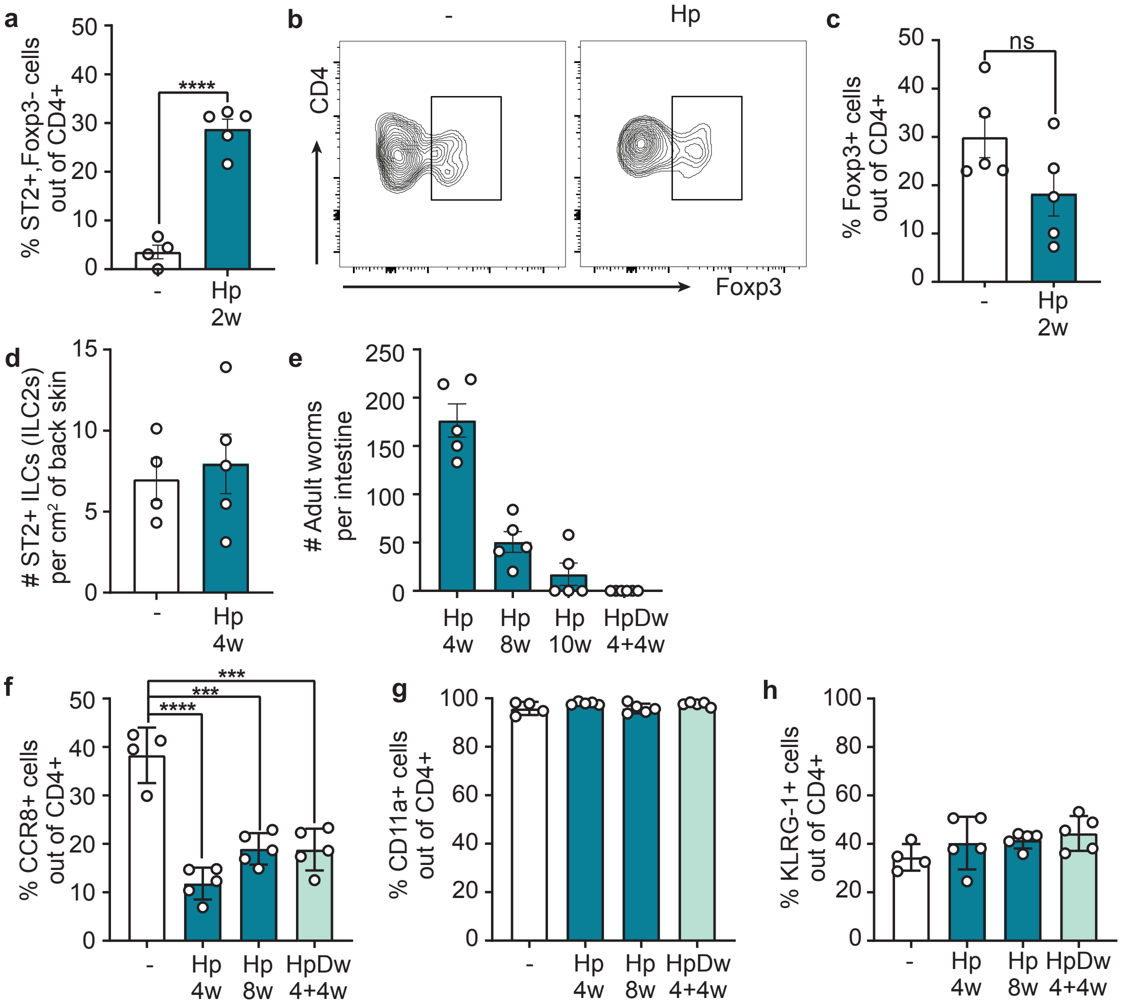


**Supplementary Figure 4: Complement to Figure 4.** Mice were infected with *H. polygyrus* (Hp) or not (-). **a** Frequency of ST2+, Foxp3- cells out of back skin CD4+ T cells. **b** Representative FACS plot of Foxp3 staining in CD4+ T cells in back skin. **c** Frequencies of Foxp3+ cells out of CD4+ back skin T cells. **d** Absolute numbers of ST2+ ILCs in back skin. **e** Adult worms counted in small intestine at indicated time points after infection with 300 L3 larvae. **f-h** Frequencies of CCR8+, CD11a+, and KLRG1+ cells out of back skin CD4+ T cells. One out of at least two independent experiments with similar results are shown. Each dot represents an individual mouse (n≥4, **a, c-h**) and bars indicate mean ± SEM. Statistical differences are depicted as ***$p < 0.001$, ****$p < 0.0001$.

### Supplementary figure 5

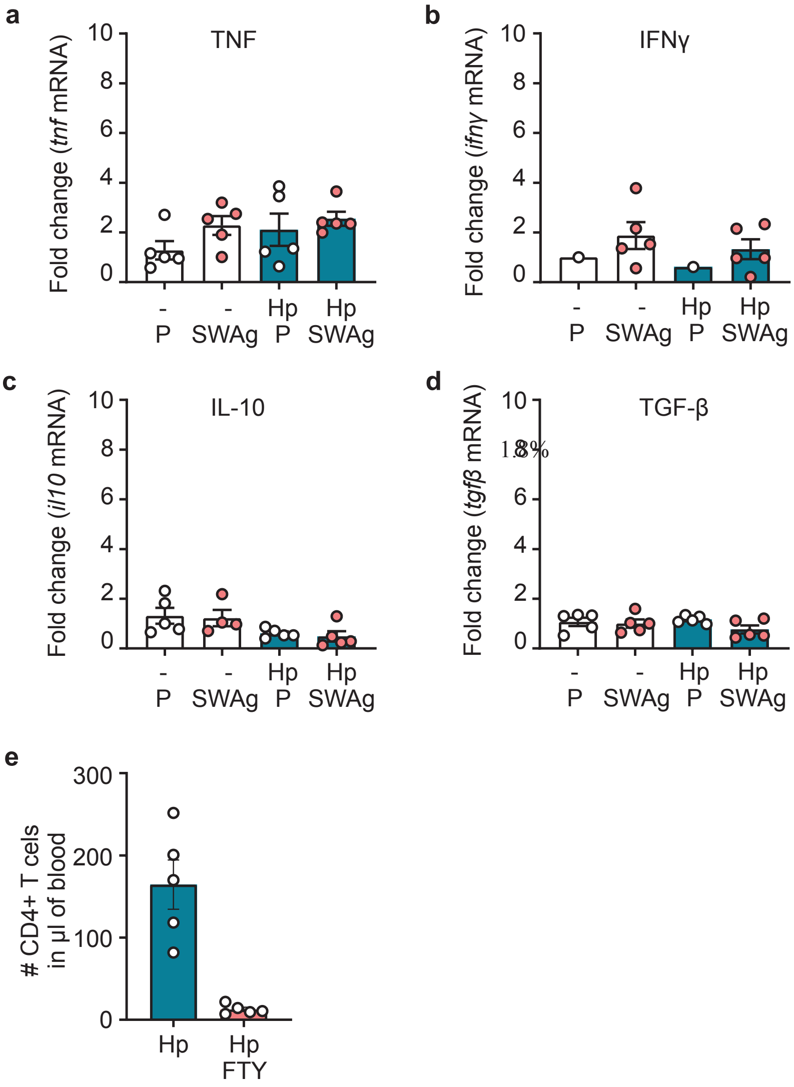


**Supplementary Figure 5: Complement to Figure 5.** Mice were infected with *H. polygyrus* (Hp) or not (-). **a-d** Soluble worm antigen from (SWAg) *H. polygyrus* (and PBS (P) in contralateral side as control) was injected in the back skin 4 weeks p.i. Fold change of mRNA expression of indicated cytokines analysed by qPCR. Five mice per group were always used, and when fewer dots are shown expression was under detection levels. **e** Number of CD4+ T cells in blood at the time of SWAg footpad injection (one day after first FTY720 treatment for Fig. 5e) counted by flow cytometry. One out of at least two independent experiments with similar results are shown. Each dot represents an individual mouse (n≥4) and bars indicate mean ± SEM.
